# Resolving a neonatal intensive care unit outbreak of methicillin-resistant *Staphylococcus aureus* to the SNV level using Oxford Nanopore simplex reads and HERRO error correction

**DOI:** 10.1101/2024.07.11.603154

**Authors:** Max Bloomfield, Sarah Bakker, Megan Burton, M. Leticia Castro, Kristin Dyet, Alexandra Eustace, Samantha Hutton, Donia Macartney-Coxson, William Taylor, Rhys T. White

## Abstract

**Objectives:** Our laboratory began prospective genomic surveillance for healthcare-associated organisms in 2022 using Oxford Nanopore Technologies (ONT) sequencing as a standalone platform. This has permitted the early detection of outbreaks but has been insufficient for single-nucleotide variant (SNV)-level analysis due to lower read accuracy than Illumina sequencing. This study aimed to determine whether Haplotype-aware ERRor cOrrection (HERRO) of ONT data could permit high-resolution comparison of outbreak isolates.

**Methods:** We used ONT simplex reads from isolates involved in a recent outbreak of methicillin-resistant *Staphylococcus aureus* (MRSA) in our neonatal unit. The raw sequence data were re-basecalled and adapter-trimmed using Dorado v0.7.0. The simplex reads then underwent HERRO correction. The resulting genome assemblies and phylogenies were compared with previous analyses (using Dorado v0.3.4, no HERRO correction and data generated by Illumina sequencing).

**Results:** Five of nine outbreak isolates were included in the analysis. The remaining four isolates had insufficient read lengths (N50 values <10,000 bp) and did not provide complete chromosome coverage after HERRO correction. The average chromosome sequencing depth for nanopore data was 147× (range: 44–220×) with an average read N50 of 12,215 bp (interquartile range (IQR): 11,439–12,711 bp). The median pairwise SNV distance between outbreak isolates from the original investigation was 51 SNVs (range: 40–68), which decreased to 3 SNVs (range: 1–15) with HERRO correction. Illumina sequencing generated a median SNV distance of 2 (range: 0–13). The resulting standalone ONT HERRO-corrected phylogeny was almost indistinguishable from the standalone Illumina-generated phylogeny.

**Conclusions:** The addition of HERRO correction meant isolates from this MRSA outbreak could be resolved to a level on par with Illumina sequencing. ONT data following HERRO correction represents a viable standalone option for high-resolution genomic analysis of hospital outbreaks, provided sufficient read lengths can be generated.

## Introduction

Next Generation Sequencing (NGS) offers huge promise for advancing surveillance of hospital pathogens, but it is still predominantly performed in larger reference or research laboratories [1]. However, this is a rapidly changing field and Oxford Nanopore Technologies (ONT) sequencing offers practical advantages as an accessible sequencing option for front-line diagnostic microbiology laboratories, including minimal footprint, low up-front capital expenses, and rapid sequence data generation [2]. Since 2022, our small-to-medium-sized laboratory lacking NGS bioinformatics expertise, has performed real-time genomic surveillance of hospital organisms using ONT sequencing to support local Infection Prevention and Control (IPC). This has been integrated into our laboratory workflow at relatively low cost and person-time expense [2] and has employed a decentralised analysis model, whereby sequencing and basic bioinformatic analysis are performed in-house and if more granular analysis is required, data are transferred to the reference laboratory for analysis by specialist bioinformaticians. This surveillance program has permitted the early detection of outbreaks, thus limiting the impact on patients and hospital services [3, 4]. Utilising publicly available sequence data, standalone ONT sequencing has been sufficient to demonstrate the clustering of outbreak isolates and permit effective IPC interventions [3, 4]. However, to date, short-read data have been required for single-nucleotide variant (SNV)-level analysis. In a prior analysis, ONT-only data resolved isolates from an outbreak of methicillin-resistant *Staphylococcus aureus* (MRSA) on our Neonatal Intensive Care Unit (NICU) to a median pairwise SNV distance of 65 SNVs, whereas Illumina sequencing produced a median distance of two SNVs due to its higher read accuracy [4]. ONT accuracy continues to improve, with updates to chemistry, basecallers and basecalling models, and more recently the introduction of Haplotype-aware ERRor cOrrection (HERRO) [5-7]. HERRO correction integrates read overlapping with artificial intelligence-based error correction of long reads which can be incorporated into ONT basecalling. Using original raw sequence data from our prior outbreak investigation, this study aimed to assess whether the addition of HERRO correction to our real-time ONT genomic surveillance system would enable SNV-level resolution of isolates.

## Methods

For this analysis, we used the originally basecalled data from White *et al*. [4], which utilised Pod5 files basecalled with Dorado v0.3.4 (https://github.com/nanoporetech/dorado, accessed 05 June 2024). Additionally, we rebasecalled Pod5s using Dorado v0.7.0. The basecalling process utilised the ‘super accuracy’ model (dna_r10.4.1_e8.2_400bps_sup@v5.0.0), with other parameters set at default. Pod5s were also rebasecalled using Dorado v0.7.0 with single-read error correction using the HERRO algorithm (implemented by the ‘dorado correct’ function using model-v1) [5]. HERRO correction recommends read lengths of >10,000 bp. Illumina sequencing data from the original investigation were also included for comparison.

NanoStat v1.6.0 and NanoQC v0.9.4 from NanoPack v1.6.0 [8] were used for initial quality assessment of the raw nanopore reads. NanoFilt v2.8.0, also from NanoPack, was used for read trimming. Initially, 50 nucleotides were trimmed from the start and end of each read to remove low-quality regions. NanoFilt was used again to filter out reads with a quality score below Q7.

Trimmed and filtered reads were *de novo* assembled and polished using the original outbreak investigation protocol for both nanopore and Illumina data [4]. However, the HERRO-corrected reads underwent *de novo* assembly using Flye v2.9.2 [9, 10] using a genome size estimate of 2.8 Mb and three polishing iterations. The corresponding Illumina reads were compared against each chromosome using Snippy v4.6.0 (https://github.com/tseemann/snippy, accessed 05 June 2024) to identify potentially erroneous variants.

We included Illumina sequence data for 13 genomes with recent evolutionary ties to the *S. aureus* sequence type (ST)97 genomes from the mid-2023 NICU outbreak [4]. These included 12 genomes from New Zealand’s national staphylococcal surveillance and one publicly available genome (Supplementary Materials, Table S1).

High-resolution analyses of genetic variants were performed using SPANDx v4.0 [11] as described previously [4]. SNV alignments were used to reconstruct phylogenies, and pairwise SNV distances were determined with snp-dist v0.6.3 (https://github.com/tseemann/snp-dists, accessed 03 July 2024). Maximum parsimony trees were reconstructed using the PAUP v4.0a heuristic search feature [12]. The resulting phylogenetic trees were visualised using FigTree v1.4.4 (http://tree.bio.ed.ac.uk/software/figtree/, accessed 03 July 2024).

## Results

Nanopore sequence data quality metrics for the nine ST97 genomes from the original outbreak are presented in Table 1. Four genomes lacked complete chromosome coverage (lower bound of median chromosome depth range includes zero) after HERRO correction due to lower read lengths and were excluded from further analysis. Figure 1 shows maximum parsimony phylogenetic trees comprising the five outbreak genomes and the 13 comparator ST97 genomes. Median pairwise SNV distances were: 2 (range: 0–13) for Illumina, 51 (range: 40–68) for ONT-only with Dorado v0.3.4, 14 (range: 9–21) for Dorado v0.7.0 without HERRO correction, and 3 (range: 1–15) for Dorado v0.7.0 with HERRO correction. HERRO-corrected and Illumina alignments were almost identical, except for two additional SNVs in the HERRO-corrected alignment and one in the Illumina alignment. Mapping Illumina reads to each chromosome revealed minimal SNV differences, mainly in insertion sequences/repeat regions and the Staphyloferrin A transporter (*sfa*A) (Supplementary Materials, Table S2).

**Table 1.**
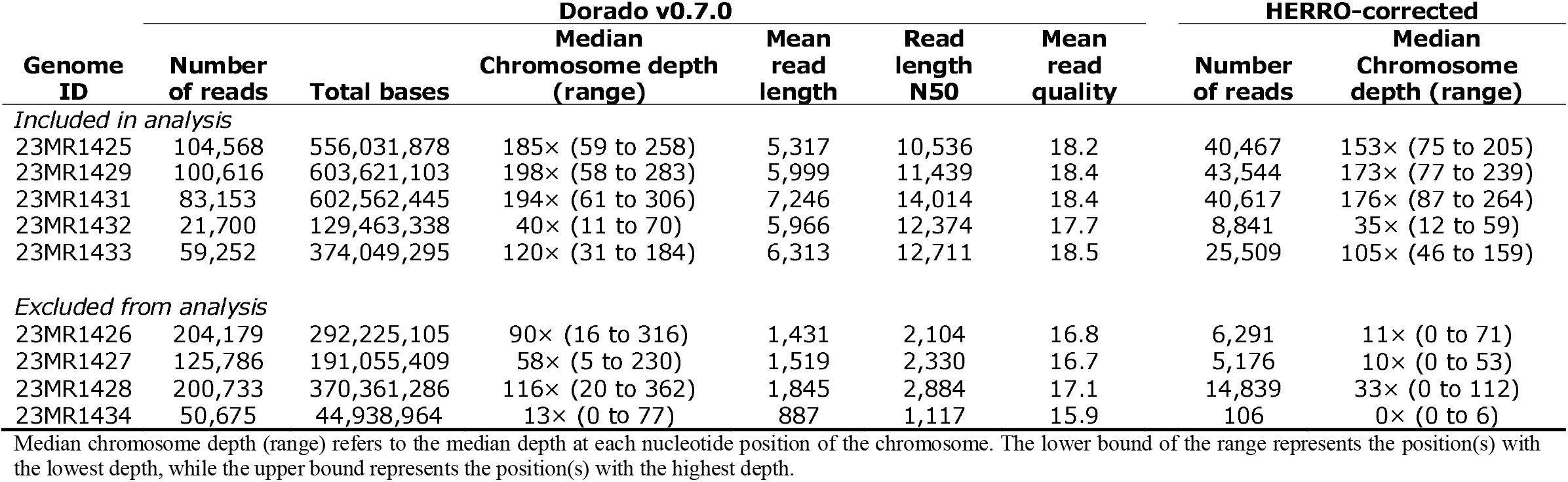
Nanopore quality control metrics for the nine Staphylococcus aureus genomes.

**Figure 1.**
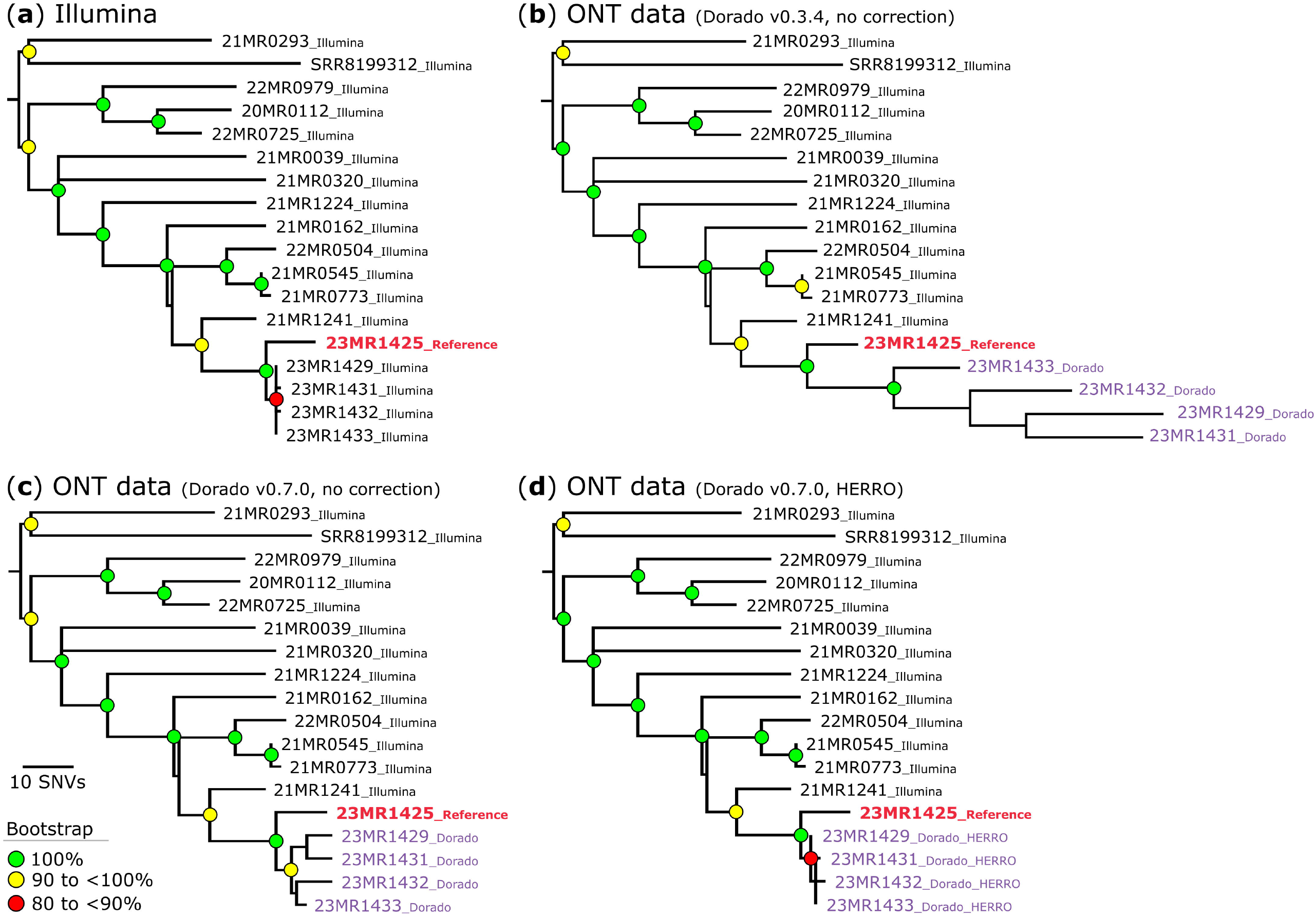
Maximum parsimony phylogeny of *Staphylococcus aureus* sequence type (ST)97. Single-nucleotide variants (SNVs) were derived from a core-genome alignment of ∼2,718,000 bp and were called against the chromosome of sample 23MR1425. SNV density filtering in SPANDx (excluded regions with ≥3 SNVs in a 10 bp window). All phylogenetic trees were rooted according to the 21MR0293 and SRR8199312 outgroup. Bootstrap values >80% (1,000 replicates) are shown. (**a**) Illumina only data for the neonatal intensive care unit (NICU) isolates. The phylogeny was inferred from 406 core-genome SNVs from 18 genomes. The consistency index for the tree was 1.0. (**b**) Nanopore only data (Dorado v0.3.4) for the NICU isolates (purple taxa). The phylogeny was inferred from 502 core-genome SNVs from 18 genomes. The consistency index for the tree is 0.95. (**c**) Nanopore only data (Dorado v0.7.0, no Haplotype-aware ERRor cOrrection (HERRO)) for the NICU isolates (purple taxa). The phylogeny was inferred from 424 core-genome SNVs from 18 genomes. The consistency index for the tree is 0.99. (**d**) Nanopore only data (Dorado v0.7.0 with HERRO correction) for the NICU isolates (purple taxa). The phylogeny was inferred from 407 core-genome SNVs from 18 genomes. The consistency index for the tree is 1.0.

## Discussion

This analysis shows that HERRO-corrected long-read nanopore data can resolve a NICU-based outbreak of MRSA to the SNV level. The assemblies and phylogenies were almost indistinguishable from those generated by Illumina sequencing. The synonymous mutation c.1045A>G in the Staphyloferrin A transporter gene (*sfa*A) in 23MR1431, 23MR1432, and 23MR1433 might be biased due to the higher chemical similarity within purines (A and G) [13, 14]. HERRO correction yielded shorter phylogenetic tree branches compared to our previous analysis [4]. Here, the median pairwise SNV distance between outbreak isolates was three, well within the proposed cutoff for relatedness of MRSA isolates of 15 SNVs [15].

These results were obtained by reanalysing the same sequence data from our original outbreak analysis, generated by our laboratory’s routine day-to-day sequencing activities. This demonstrates the viability of using ONT sequencing for real-time, high-resolution outbreak detection in hospitals, making it suitable for front-line clinical microbiology laboratories. ONT sequencing offers low up-front capital costs, a minimal physical footprint, and long reads that enable complete *de novo* genome assembly, as well as the detection of mobile genetic elements and plasmids, crucial for IPC and antimicrobial resistance gene detection.

A limitation of this analysis was that only a small number of isolates from a single species were included. To confirm these results, analysis of larger datasets and investigations of the effects of differing depths of genome coverage on HERRO-corrected assemblies are required. Of note, our lowest depth isolate had a median chromosome depth of 35×, with the lowest region being 12× (Table 1). Despite this, an Illumina-equivalent assembly was generated. However, the small number of available isolates was further reduced by excluding four genomes due to insufficient read lengths. This highlights a potential limitation of the implementation of the HERRO correction approach by ONT for their solo long reads, particularly for organisms that typically produce shorter reads, which has been our experience with pathogens such as *Clostridioides difficile* [2, 3].

Recent publications highlight advances in ONT sequencing accuracy through updated chemistry, duplex reads, and ‘methylation-aware’ basecallers, which have significantly improved ONT-only phylogenomic analyses and reduces the need for hybrid genome assembly with short read data [6, 7]. Our analysis further supports ONT sequencing as a viable standalone option for high-resolution surveillance of hospital pathogens, with HERRO correction resulting in highly accurate assemblies.

## Acknowledgements

We acknowledge the facilities, and the scientific and technical assistance of staff at Awanui Labs, formerly known as Southern Community Laboratories (SCL) (Wellington, New Zealand). We are grateful to the diagnostic microbiology laboratories across New Zealand for contributing isolates and associated data to the surveillance programmes at the Institute of Environmental Science and Research (ESR). We thank the Next Generation Sequencing Team and the Antimicrobial Resistance Laboratory at ESR (Porirua, New Zealand) for their valuable scientific and technical assistance. We acknowledge and thank the following ESR staff for their valuable feedback: Xiaoyun Ren (Kenepuru Science Centre) and Vinko Besic (Kenepuru Science Centre). This research was made possible by the Computational Science Team and the High-Performance Compute (HPC) platform at ESR, with specific thanks to Russell Smithies and Shane Sturrock for HPC support and helpful discussions about software pipelines. Finally, we thank the New Zealand Ministry of Health/Manatū Hauora for funding this work.

## Data availability statement

The study sequences are available in the National Center for Biotechnology Information (NCBI) BioProject accession number PRJNA1046639. The raw sequence read data from Illumina, and Oxford Nanopore Technologies (ONT) basecalled with Dorado v0.3.4, have been deposited to the NCBI Sequence Read Archive (SRA) under accession numbers SRR26992863 to SRR26992891. The re-basecalled (Dorado v0.7.0) ONT raw sequence read data generated in this study are available under accession numbers SRR29788345 to SRR29788353. The complete assembly for strain 23MR1425 has been deposited to GenBank under the accession numbers CP143800 and CP143801.

## Contribution

*Writing – Original Draft*: M.Bl. and R.T.W; *Writing – Review & Editing*: M.Bl., S.B., M.bu., M.L.C., K.D., A.E., S.H., D.MC., W.T., and R.T.W.; *Conceptualization*: M.Bl. and R.T.W.; *Investigation*: M.Bl. and R.T.W.; *Methodology*: S.B., M.bu., M.L.C., K.D., A.E., S.H., and R.T.W.; *Data Curation*: R.T.W.; *Visualization*: R.T.W.; *Formal Analysis*: R.T.W.; *Project Administration*: M.Bl. and R.T.W.; *Funding Acquisition*: M.Bl. and D.MC. All authors have read and agreed to the published version of the manuscript.

